# Deep learning-enhanced single-molecule spectrum imaging

**DOI:** 10.1101/2023.05.08.539787

**Authors:** Hao Sha, Haoyang Li, Yongbing Zhang, Shangguo Hou

## Abstract

Fluorescence is widely used in biological imaging and biosensing. Rich information can be revealed from the fluorescence spectrum of fluorescent molecules, such as pH, viscosity and polarity of the molecule’s environment, and distance between two FRET molecules. However, constructing the fluorescence spectrum of a single fluorescent molecule typically requires a significant number of photons, which can suffer from photobleaching and therefore limit its potential applications. Here we propose a deep learning-enhanced single-molecule spectrum imaging method (SpecGAN) for improving the single-molecule spectrum imaging efficiency. In SpecGAN, the photon flux required to extract a single-molecule fluorescence spectrum can be reduced by 100 times, which enables it two orders of magnitude higher temporal resolution compared to the conventional single-molecule spectrometer. The concept of SpecGAN was validated through numerical simulation and single Nile Red molecule spectrum imaging on support lipid bilayers (SLBs). With SpecGAN, the super-resolution spectrum image of the COS-7 membrane can be reconstructed with merely 12,000 frames of single-molecule localization images, which is almost half of the previously reported frame count for spectrally resolved super-resolution imaging. The low photon flux requirement and high temporal resolution of SpecGAN make it a promising tool for investigating the molecular spectrum dynamics related to biological functions or biomolecule interactions.

## 1. Introduction

Since the invention of optical microscopy, it has become an essential tool in life science research due to its non-invasive nature and specificity. Over the last several decades, the emerging super-resolution fluorescence microscopy overcome limits of optical diffraction and pushed the spatial resolution up to several nanometers^1-5^, which can reveal unprecedented fine structure detail of organelles in cells. In fluorescence microscopy, the fluorescence intensity, i.e., fluorescence photon emission rate is usually used to build a morphology image of the sample. Aside from the intensity information, fluorescence signal also has many other characteristics, such as fluorescence spectrum^6^, fluorescence lifetime^7^, and fluorescence polarization^8^. These characteristics indicate the microenvironment biophysical properties of the fluorescent molecules, such as pH, viscosity, and polarity.

In recent years various spectrum imaging methods were developed. Fluorescence spectrum can provide additional information that cannot be acquired by intensity imaging, such as molecular structure^9, 10^, membrane potential^11^, spatial arrangement of biomolecules^12, 13^, and the microenvironment surrounding molecules^14-16^. Spectrum imaging has been applied to various research fields. For instance with fluorescence spectrum imaging, the stereoisomers of borondipyrromethene (BODIPY) can be well resolved and their photophysical properties and excitation dynamics can be further investigated^17^. Combining the pH-sensitive photochromic dye and spectrum imaging is able to disclose the alternative route of nanoparticles reaching lysosomes and to observe the pH changes in real-time^18^. The surface hydrophobicity of α-synuclein (αS), a Parkinson’s disease related protein, can also be characterized with the fluorescence spectrum of solvatochromic dye^19^. Usually, the fluorescence spectrum is measured by fluorescence spectrophotometer. However, the fluorescence spectrophotometer is not applicable to cellular experiment, which precludes its further cellular applications.

To acquire the morphological and spectral information of cells simultaneously, spectrally resolved super-resolution microscopy (SR-SRM) has been proposed in recent years^15, 20, 21^. This method has been applied to reveal the nanoscale spectral features of different dyes on cell membrane^15, 21^. Besides, the polarity heterogeneity of the plasma membrane and organelles membranes with different cholesterol levels were revealed with Nile Red labeling^22^. On the other hand, owing to the limited fluorescence emission rate and total emission photons of a single molecule, obtaining a highly precise fluorescence spectrum of a single molecule in the short collection time period is challenging. Particularly in intracellular imaging, the presence of autofluorescence further deteriorate the spectrum imaging. To obtain an accurate fluorescence spectrum, multiple spectra are usually collected and averaged. For SR-SRM imaging of mammalian cells, reconstructing a spectral mapping image typically requires 15,000-30,000 frames of localization image, resulting in a poor temporal resolution and precluding the spectral dynamics analysis^23^.

Deep learning technology has become increasingly popular in the realm of microscopic imaging, with a broad range of applications including denoising^7, 24^, modal conversion^25, 26^, resolution enhancement^27^, downstream task analysis^28, 29^, and high-speed imaging^30-32^. In single-molecule spectral imaging, deep learning has also been employed to address the problem of spectral noise. Convolution neural networks (CNNs) have been found to be highly effective in feature extraction, enabling the accurate classification of three-color labeled cell samples with low rates of molecular misidentification^33^. However, this approach was specifically designed for classifying different dyes and cannot be applied to identify the continuous spectral changes in different environments. In image generation tasks, the generative adversarial network (GAN)^34^ has emerged as a popular generative model. While originally proposed for unsupervised learning, GAN has been shown to be effective in supervised tasks, such as super-resolution and semantic segmentation^35, 36^.

In this paper, we propose a novel deep learning-based single-molecule spectrum imaging method called SpecGAN, which is tailored to denoise single-molecule spectra and accurately identify the spectral characteristics of environment-sensitive fluorescent dyes. The SpecGAN is validated with simulation, support lipid bilayers (SLBs) imaging and cell membrane superresolution spectrum imaging. Notably, implementing SpecGAN does not require modifications to the current spectrally resolved super-resolution microscopy (SR-SRM) system, and it holds promising potential for real-time investigations of single-molecule spectral dynamics owing to its low photon flux requirement to construct the spectrum.

## 2. Results

### 2.1 Workflow of SpecGAN

Generative Adversarial Networks (GANs) consist of two main components: a generator and a discriminator. The generator is responsible for creating synthetic data instances by learning the underlying distribution of the real dataset while the discriminator, on the other hand, is a neural network designed to distinguish between real and synthetic data instances. Similar to the GAN model, SpecGAN contains a generator and discriminator. The generator of SpecGAN takes the 1-d spectral data with noise into a UNet-based network and outputs clean spectrum data. The UNet is a convolutional neural network (CNN) architecture designed for semantic segmentation tasks. The network architecture consists of a contracting path that captures context and a symmetric expanding path that enables pixel-to-pixel mapping^37^. In UNet, the encoder uses convolution layers and maximum pooling layers for feature extraction, while the decoder uses binary linear interpolation and convolution layers for upsampling and information recovery, respectively. For the discriminator of SpecGAN, auxiliary classification tasks^38^ are adopted to enhance the model’s focus on environmental information. In addition to determining whether the spectral curve is generated from the generator, the discriminator is also responsible for classifying the environments represented by the spectrum. More details about SpecGAN can be found in the method section.

To validate the performance of SpecGAN, we built a single-molecule spectrum imaging microscopy (Fig. 1a). The fluorescence is directed to two paths by a beam splitter. The fluorescence in path 1 is used for recording the location information of molecules while path 2 is used for recording the fluorescence spectrum. An example of fluorescent bead spectrum imaging is shown in Fig.1(a). As previously mentioned, to obtain the fluorescence spectrum of single molecules, it is necessary to average multiple spectral data from the same environment to counteract the influence of noise. As a demonstration, we simulated the average fluorescence spectrum with a different amount of fluorescent spectrum data (Fig.1 (b)). It can be seen that several hundred single-molecule spectra data are required to produce a well-resolved averaged spectrum. In this study, we choose Nile Red as a fluorescence probe to evaluate the performance of SpecGAN. Nile Red is a widely used fluorescent marker to monitor the changes of polarity in spectrally resolved super-resolution imaging because it exhibits a significant blue shift in the lipid order (Lo) phase compared to the lipid disorder (Ld) phase^39^, as shown in Fig.1(c).

**Fig. 1.**
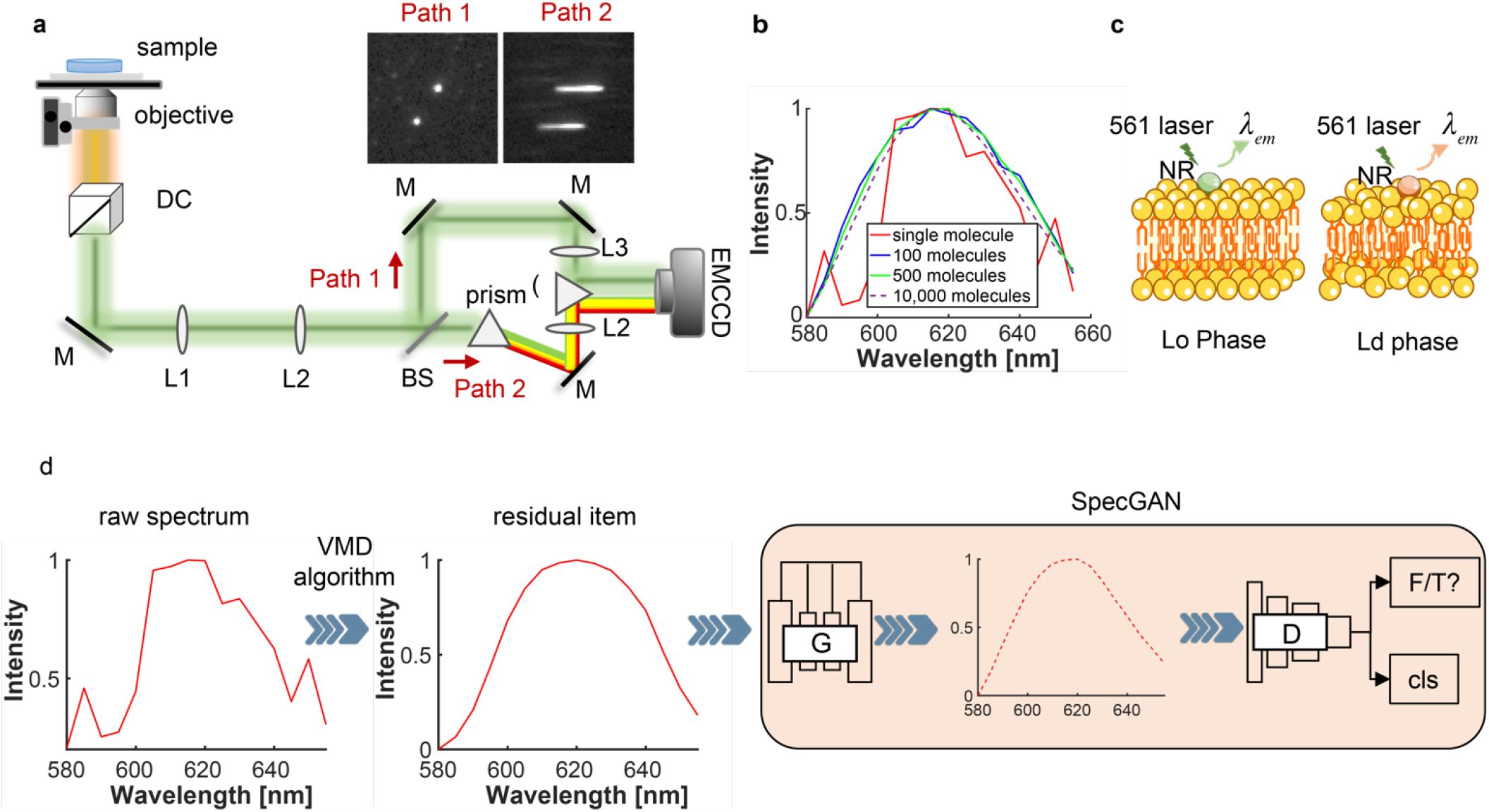
Experimental setup and data process overview. (a)Schematic diagram of the single-molecule spectrum imaging microscopy setup. The inset shows the spectra and position images of the fluorescent bead. (b) Averaged spectra with different mounts of recorded molecules. (c) Schematic of Nile Red in different lipid environments. The emission spectra peak of Nile Red (NR) changes with the lipid orders. Lo: lipid order; Ld: lipid disorder. (d) Overview of the SpecGAN structure, where the residual item is extracted from the raw spectrum by the VMD algorithm, and the SpecGAN outputs the denoised spectral curve of the fed residual one.

The flowchart of spectrum imaging with SpecGAN is illustrated in Fig.1(d). Due to the low signal-noise ratio (SNR) of the raw spectrum of a single molecule, it is challenging for the SpecGAN model to reach stabilization gradually during training. Thus, previous to feed the raw spectral data into SpecGAN, the variational modal decomposition (VMD) algorithm is used to adaptively decompose a spectral curve into several intrinsic mode functions (IMFs) and a residual item^40^. The residual item of VMD, which contains the inherent feature of the raw spectral curve and is always smooth due to the removal of high-frequency noise components, is used as input for the SpecGAN. For the construction of training datasets, the average of all spectral curves is taken as the ground truth. This workflow aims to generate a fluorescence spectrum similar to the ground truth spectrum using the fewest single-molecule spectral data.

### 2.2 SpecGAN performance evaluation with simulated data

The performance of SpecGAN is firstly tested with simulation data based on the spectral shifts of Nile Red in different solvents^41^. The Nile Red spectra in different solvents (Acetone, CHCl3, C3H8O, MeOH, and EtOH) are measured with fluorescence spectrophotometer (Lumina, Thermo Scientific) and used as the ground truths during the training stage (Fig. 2a). The fluorescence centroid in different solvents show different spectral characters, with the blue shift being most pronounced in chloroform and the redshift in MeOH (Fig. 2b). Additionally, these solvent categories are further labeled as the environmental classes for the supervision of the auxiliary discriminator. To generate the spectrum with noise, these ground truths are added with Gaussian, Poisson, and salt & pepper noise. The simulation dataset comprises approximately 50,000 spectral curves with varying noise levels and spectral centroids. This dataset is divided into training, validation, and test subsets, maintaining a ratio of 0.8, 0.1, and 0.1, respectively.

**FIG. 2.**
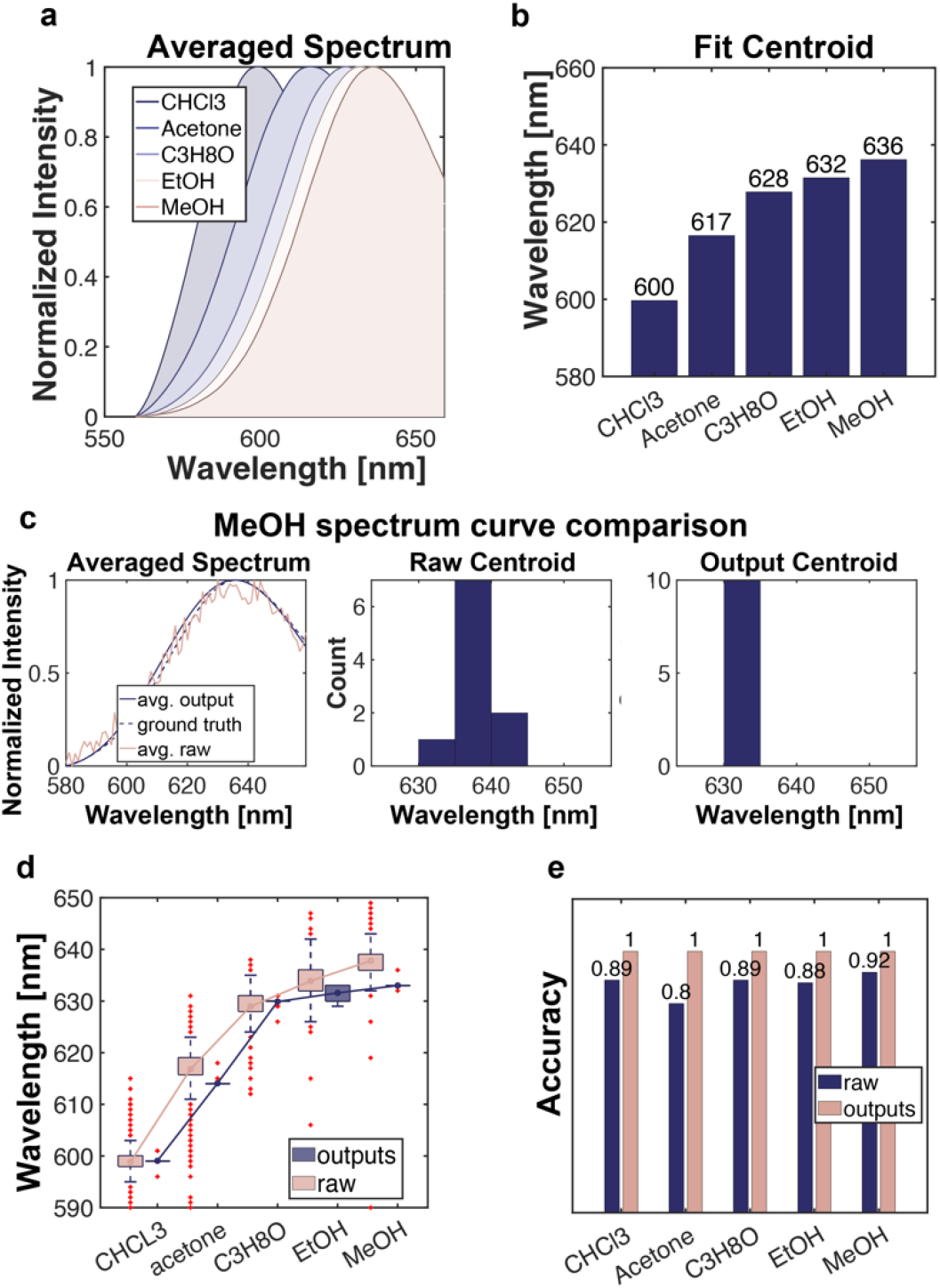
Performance evaluation of SpecGAN with simulated data. (a) Spectra of Nile Red in different solvents measured by fluorescence spectrophotometer. (b) The spectral centroids of Nile Red spectra in different solvents. (c) Comparison of raw spectrum with noise and the spectrum output from SpecGAN. Solvent: MeOH. (d) Fluorescence spectrum centroid of Nile Red in different solvents calculated with raw spectra and SpecGAN outputs. (e) Fluorescence spectrum maxima identification accuracy comparison.

As shown in Fig.2c, the averaged spectrum of 10 raw fluorescence spectra significantly deviated from the ground truth and the centroid histogram is dispersed. This result indicates that the random noise in the simulation data may interfere with the analysis of the molecular fluorescence spectrum, resulting in an incorrect spectral centroid measurement. In contrast, the average spectrum of SpecGAN outputs is closer to the ground truth, and the centroid of the spectrum can be easily identified. Correspondingly, the outputs of SpecGAN show smaller variances, indicating that SpecGAN can give a centroid prediction with high confidence (Fig. 2d). We then compared the fluorescence spectrum maxima identification accuracy of SpecGAN output data and raw data. We found that the identified centroids with SpecGAN nearly 100% deviate less than 5 nm from the ground truth. Compared with raw data, SpecGAN output data show a significant accuracy improvement, specifically, with a maximum of 20% accuracy improvement for the spectra in acetone solvent measurement.

### 2.3 Results on SLBs data

To demonstrate the biological compatibility of SpecGAN, the SLBs spectrum imaging was conducted. SLBs are artificially constructed, two-dimensional lipid structures that mimic the properties of natural cell membranes. SLBs serve as valuable tools for studying the biophysical and biochemical properties of cell membranes, as well as for investigating processes such as membrane-protein interactions, lipid organization, and the dynamics of membrane components^8^. We made the SLBs by mixing the lipids 1,2-dioleoyl-sn-glycero-3-phosphocholine (DOPC), sphingomyelin (SM), and cholesterol (Chol) in chloroform in different proportions. These proportions were selected to induce the emission spectrum shift of Nile Red. We utilized five distinct mixing ratios of DOPC, SM, and Chol in this study, specifically 1:0:1, 1:1:0, 1:1:1, 1:3:1, and 0:1:1, representing the respective proportions of the DOPC, SM, and Chol components. These mixtures are abbreviated as DC, DS, DSC, DSC311, and SC, respectively. To prepare the SLBs, we employed the bicelle adsorption and fusion method, which facilitates the formation of stable bilayer structures on glass surfaces^42^.

In this work, the spectra of Nile Red in five different SLBs compositions were collected using single-molecule spectrum imaging microscopy. Approximately 60,000 single-molecule spectral data for each category are averaged to create the ground truths (Fig. 3a, Fig. S6). For the SLBs Nile Red spectra dataset recorded by single-molecule spectrum imaging microscopy, we found that the noise has a non-negligible effect on signal processing, which may make the deep learning model hard to converge during training. Therefore, it is necessary to exclude the spectral data with significant noise. In the training of SpecGAN, the SLBs dataset is manually pre-screened to exclude poor-quality spectra data. A signal is considered usable if the difference between the centroid of the residual item and the average spectrum is smaller than ±15nm, or if the root mean square error (RMSE) is smaller than 0.3. Compared with raw spectra data, the fluorescence spectral centroid localization accuracy of SpecGAN outputs is improved by up to 62% and the variance of the SpecGAN outputs is reduced by about 4 times. (Fig.S6 - Fig. S8). Moreover, using SpecGAN, the Nile Red spectrum can be reconstructed with just 10 molecular spectral data points, whereas approximately 500 to 1000 molecules were necessary for reconstructing a similar Nile Red spectrum in raw data. This indicates that SpecGAN can potentially reduce the photon flux requirements by up to 100 times.

**FIG. 3.**
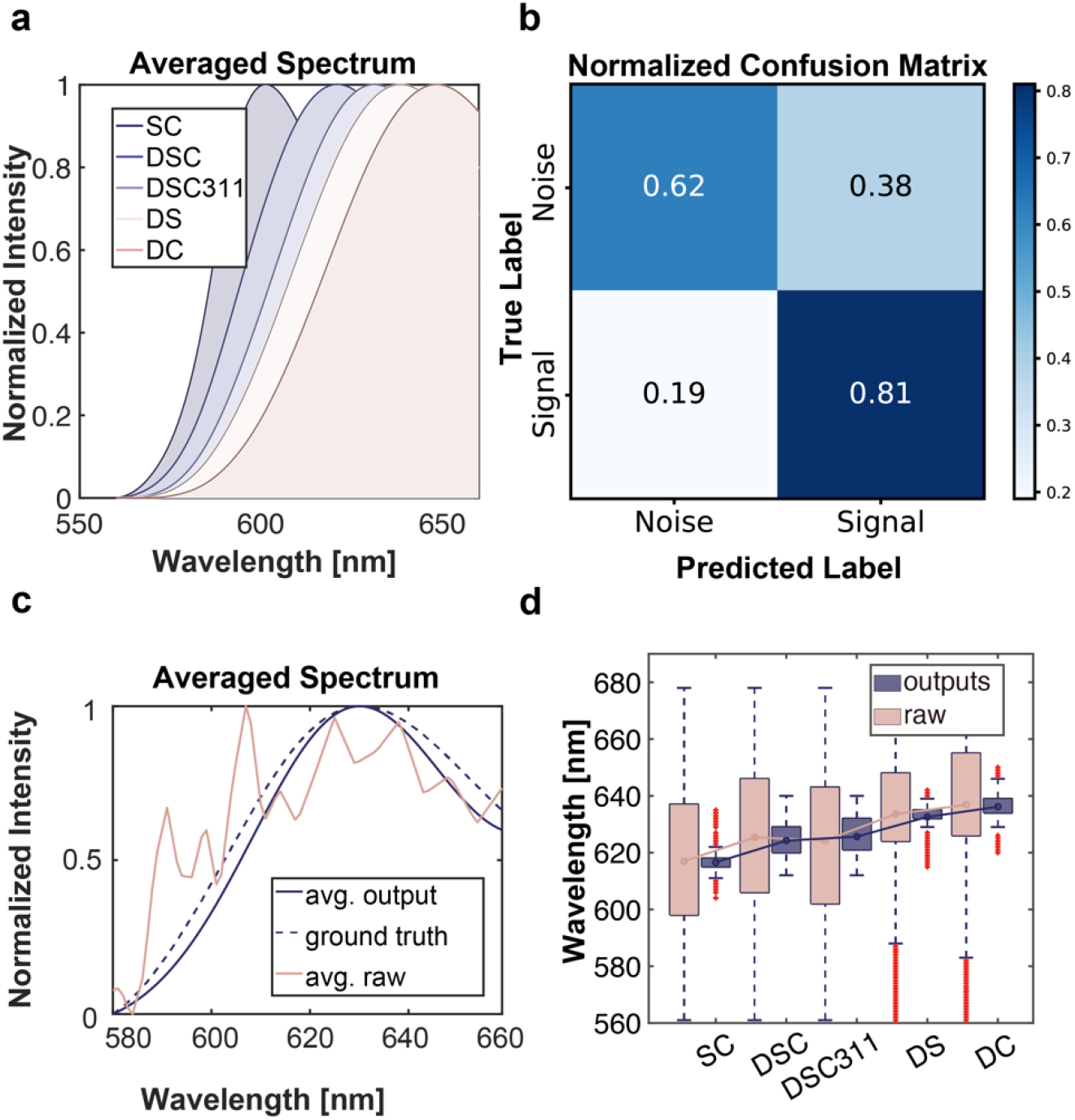
Performance evaluation of SpecGAN with SLBs data. (a) Averaged spectra of Nile Red in SLBs with different components acquired by single-molecule spectrum imaging microscopy. (b) Normalized confusion matrix of ResNet-based classification model. (c) The comparison between raw spectrum and the output of SpecGAN in DOPC:SM:Chol (3:1:1). (d) Boxplot of centroid calculated by raw spectra and SpecGAN outputs.

SpecGAN has demonstrated strong denoising capabilities when applied to manually screened data. To automate the data screening process, we trained a ResNet-based classifier as a replacement. This classifier can automatically determine the signal quality, and the screened data is then fed to the SpecGAN, which has already been trained on manually screened datasets. The normalized confusion matrix of the ResNet-based classification model is displayed in Fig. 3b. After 150 training iterations, the model achieves a precision and recall rate of 68% and 81%, respectively. Then we test its denoising capabilities with 10 single-molecule spectra data (Fig. 3c, 3d). In the raw SLBs dataset, the centroid cannot be identified from the limited numbers of single molecule spectrum, and the variance is much higher than the SpecGAN outputs. Moreover, the wavelength of DSC centroid shows a slight red shift compared to DSC311, which is not consistent with the ground truth. As a comparison, SpecGAN achieves 81% identification accuracy, showing an improvement of 56% over the raw data analysis (Fig. S10). It should be noted that the centroid determined by SpecGAN for the SC category deviates slightly from the average fluorescence spectrum, but still has a lower variance compared with raw data. The deviation may be caused by the poor performance of the classifier in the SC dataset.

### 2.4 SpecGAN enhanced super-resolution spectrum imaging

The super-resolution spectrum imaging technique or spectrally resolved stochastic optical reconstruction microscopy (SR-STORM) has been developed in recent years^15, 20, 21^. Incorporating spectral information into super-resolution imaging not only adds an additional dimension for analysis but also enhances localization precision. The SpecGAN has the capability to further improve superresolution spectrum imaging. Here we performed a super-resolution spectrum imaging of Nile Red labeled COS-7 cells to demonstrate the performance of SpecGAN. Compared to super-resolution spectral imaging reconstructed using raw data (raw SR-STORM), SpecGAN significantly improves spectral precision (Fig. 4a,b). The spectral distribution in COS-7 cells appears discontinuous and noisy for raw SR-STORM (Fig. 4a), while the SpecGAN output image displays a more continuous and concentrated spectral distribution (Fig. 4b). Additionally, we compared the spectrum of a pixel for raw SR-STORM and SpecGAN output and observed that the spectrum in raw SR-STORM is heavily affected by noise, ultimately impacting the accuracy of spectrum maxima localization (Fig. 4c). In contrast, the SpecGAN output yields a smoother spectrum curve, facilitating more precise spectrum maxima localization. In a set of 12,000 localization images, the majority of pixels have fewer than 10 localizations (Fig. 4d). As a result, addressing noise interference is crucial for fast spectrally resolved super-resolution imaging. Herein SpecGAN provides a novel approach to analyze spectral features at the single-molecule level.

**FIG. 4.**
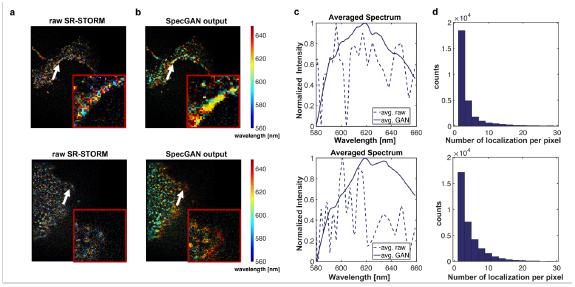
SpecGAN enhanced superresolution spectrum imaging. (a) Super-resolution spectrum imaging with raw-STORM. (b) Super-resolution spectrum imaging with SpecGAN. Each detected single molecule is color-coded according to its spectral centroid. The inset is a magnified view of the region indicated by the white arrow. (c) Comparison of averaged single-molecule spectral curve at the cell membrane pointed by the arrow in (a) and (b). (d) Histogram of numbers of localization per pixel. Total frames: 12,000.

## 3. Discussion and Conclusion

In this study, we presented a novel deep learning-enhanced single-molecule fluorescence spectrum imaging technique called SpecGAN, which allows for the effective extraction of single molecule spectra with significantly improved signal-to-noise ratio and accuracy. The SpecGAN was applied to spectral imaging of Nile Red labeled SLBs and super-resolution spectrum imaging of Nile Red labeled COS-7 cells with low photon flux. SpecGAN effectively reduces noise interference in spectrum construction, resulting in fewer fluorescence photons required to construct the spectrum. Compared to conventional spectral super-resolution imaging methods, SpecGAN demonstrates approximately a 2-fold improvement in temporal resolution. In the future, SpecGAN could be combined with single-molecule tracking to monitor real-time biochemical interaction processes, such as changes in the tumor microenvironment during drug delivery^43, 44^. Overall, this work introduces a promising approach for fast spectrum imaging for weak fluorescence signals, which holds significant potential for a wide range of applications in the study of biological interaction dynamics.

## Methods

### Pixel shifts and wavelength mapping

A bandpass filter (FF01-591/6-25, Semrock) is used to collect at least 6 paired points for calculating the mapping matrix. The code implementation is based on Xu *et al*^45^. The mapping matrix can transform the pixel coordinates in the position channel to the position of 591 nm in the spectral channel. Further, to establish the mapping relationship between the pixel shifts and wavelength, fluorescent beads (BangsLab FSDG003 and FSSY002) are imaged and the pixel coordinates of three specific wavelengths in the spectral channel are recorded using bandpass filters (FF01-532/3-25, LL02-561-12.5, LL01-638-12.5, Semrock). The pixel-wavelength calibrated curve is fitted by a third-order polynomial finally, as shown in Fig.S3 and Tab.S1. The center coordinates of single molecules in each frame are detected using ThunderSTORM plugin^46^. Therefore, the spectral curve of each molecule and the average value which is considered the ground truth can be obtained.

### The VMD-based pre-processing

In order to enhance the performance of our model by reducing the impact of noise, we propose a data preprocessing method based on VMD. As a signal process method without prior knowledge, VMD has been applied to various signal tasks^47^. It assumes that a signal is composed of a series of sub-signals with specific center frequencies and limited bandwidths. The solution to this variational problem involves using the Weiner filter and the Hilbert transform^40^. Utilizing the VMD algorithm, the spectral information corrupted by noise can be separated into different IMFs and a residual item. The residual signal is relatively smoother and partially removes the interference of high-frequency noise. Thus, we use the residual component derived from VMD as the input of the SpecGAN model, which can lead to superior performance. More details about the VMD algorithm can be found on SI.

### ResNet-based data screening

In this work, a two-stage network is proposed to identify the accurate centroid of the spectrum collected by the SR-SRM system at the single-molecule level. The structure of the SpecGAN is shown in Fig.5. The presence of environmental noise results in low SNR in single-molecule spectrum images, which precludes the precise single-molecule spectral features analysis. To address this issue, a classifier, as depicted in Fig.5(a), is proposed to exclude signals with too low SNR from the SpecGAN training and testing process. The traditional 2-D convolution layer is adapted to the 1-D spectral data by converting the number of parameters from 120 to 4096 using a fully connected network and reshaping it to 64×64×1. Subsequently, feature extraction is performed through a simplified ResNet^48^. The output of the classification model is a binary result indicating whether the current signal is useful or not. This model is trained using a binary cross-entropy loss function.

**FIG.5.**
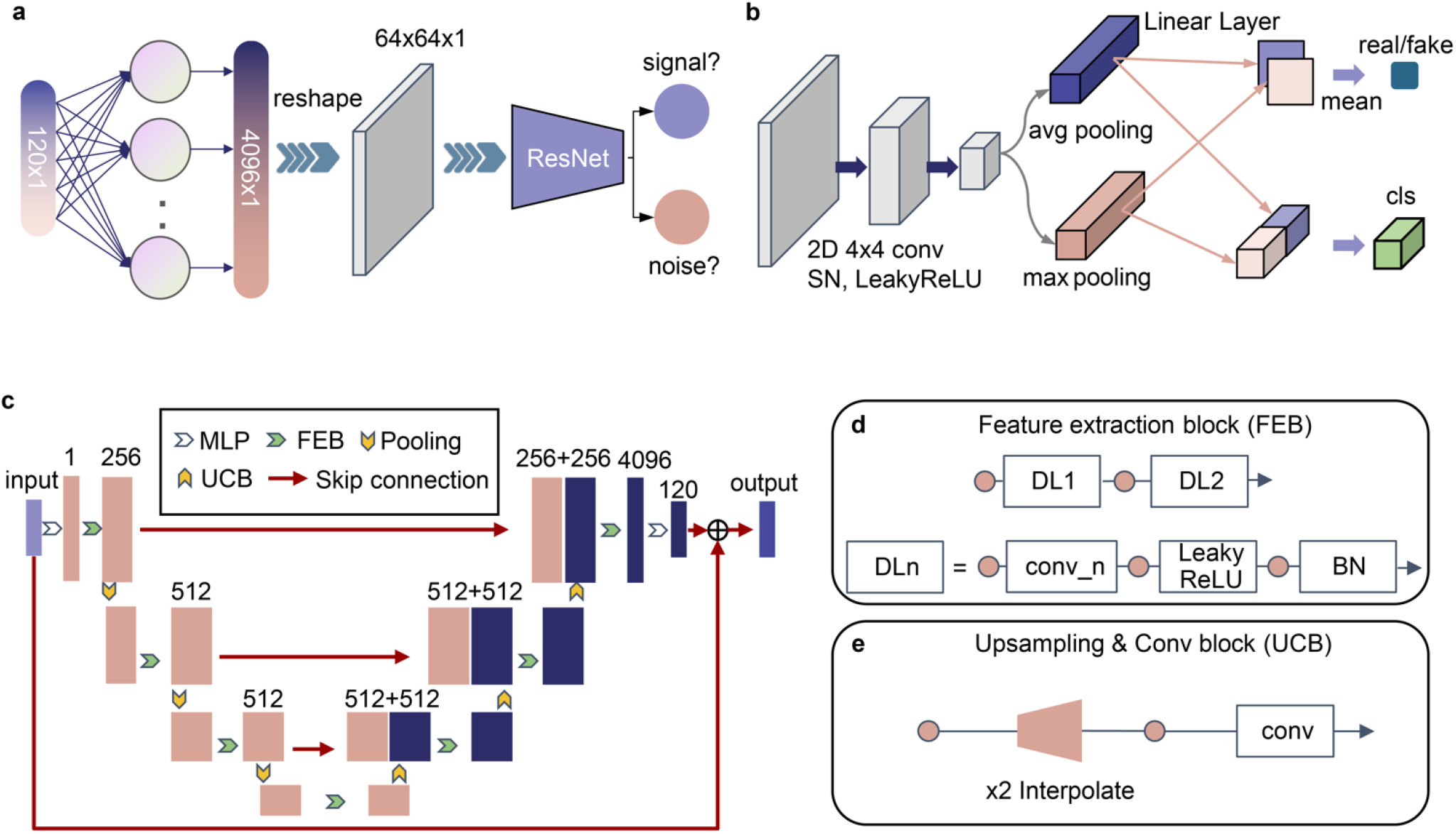
Workflow of SpecGAN. (a) A two-stage network model for filtering low-SNR spectral signals. The ResNet-based classification model is employed to judge whether the inputs are valid signals or noise and only the valid signals are fed to the subsequent SpecGAN training and testing in SLBs and cell data. (b) Discriminator of SpecGAN with the task of determining whether the data is real or fake and the corresponding environmental category of the current Nile Red molecular spectrum. (c) Generator of SpecGAN based on a simplified UNet architecture, with details of the feature extraction block (FEB) and the upsampling & convolution block (UCB) shown in (d) and (e). The FEB extracts abstract features and maps them to a lower-dimensional representation, while the UCB uses a bilinear interpolation layer to upsample the lower-dimensional representation.

### Details of the SpecGAN

SpecGAN aims to produce a pure spectral curve that can identify environmental features from the VMD residual component of the raw signal. It contains a discriminator and a generator. The generator in Fig.5(c) is based on the UNet architecture and is composed of 8 sub-blocks. The encoder uses convolution layers and maximum pooling layers for feature extraction, while the decoder uses binary linear interpolation and convolution layers for upsampling and information recovery. The corresponding sub-blocks between the encoder and decoder are concatenated for information interaction. The output of the UNet is then passed through a linear layer to ensure that it has the same dimensions as the input spectral data and then added to the original input to produce the final denoised spectral curve. In particular, the LeakyReLu activation layer and batch normalization (BN) layer are utilized in the feature extraction block (FEB).

In the discriminator depicted in Fig.5(b), auxiliary classification tasks are designed to further improve the performance. The discriminator needs to determine the confidence of the pure spectral curve and classify the environments represented by it. The labels have been previously described in detail. For other environment-sensitive dye molecules, changes in environmental conditions such as pH, viscosity, and micro-environment can also be encoded as labels. The discriminator initially extracts features using three convolution layers with the spectral normalization (SN) method^49^. Subsequently, a global average pooling and a global maximum pooling layer reduce the feature map to a size of 1×1×128. Finally, the downscaled pooling features are processed using linear layers based on the requirements, and the corresponding averages are used for auxiliary classification or GAN tasks.

### Loss function

In terms of the loss function of SpecGAN, we use Wasserstein GAN loss^50^ to improve the stability during training, while the L1 loss is introduced to determine the distance between the input spectrum and the average spectrum. The auxiliary classification loss is employed to measure the accuracy of category classification. Additionally, the 1-D total variation (TV) regular term loss function is included to make promote output spectral curves smoother. During training, the generator and discriminator parameters are updated in an alternating manner.

## SUPPLEMENTARY MATERIAL

See supporting information for more results and details of experiments.

## Conflict of interest

The authors declare no conflict of interest.

## Data Availability Statement

The code implementation and data for training and testing in this work are available at our GitHub repository, git@github.com:hitsh95/SpecGAN.git. The raw images acquired by SR-SRM system are available from the corresponding authors upon reasonable request.

## ACKNOWLEDGMENTS

S. Hou would like to acknowledge support from National Natural Science Foundation of China (22204106), the Guangdong Basic and Applied Basic Research Foundation (2021A1515110710) and the Guangdong Pearl River Talents Program (2021QN02Z631). Y. Zhang would like to acknowledge support from National Natural Science Foundation of China (62031023), Shenzhen Science and Technology Project (JCYJ20200109142808034 & GXWD20220818170353009 & JSGG20191129110812708), and Guangdong Special Support (2019TX05X187). S. Hou and S. Hao wish to thank Dr. Yan Zhao in Shenzhen Bay Laboratory for her help on fluorescence spectrophotometer and thank Prof. Chenguang Wang in Jilin University for the helpful discussion. S. Hou and S. Hao also would like to express appreciation to Prof. Xu at the University of California, Berkeley for the discussion on the alignment of the spectral and position channels.

## Notes

### Competing Interest Statement

The authors have declared no competing interest.

## REFERENCES

1. M. J. Rust, M. Bates, and X. Zhuang, “Sub-diffraction-limit imaging by stochastic optical reconstruction microscopy (STORM),” Nat Methods 3, 793–795 (2006).

2. A. Sharonov, and R. M. Hochstrasser, “Wide-field subdiffraction imaging by accumulated binding of diffusing probes,” Proc Natl Acad Sci U S A 103, 18911–18916 (2006).

3. M. D. Lew, S. F. Lee, J. L. Ptacin, M. K. Lee, R. J. Twieg, L. Shapiro, and W. E. Moerner, “Three-dimensional superresolution colocalization of intracellular protein superstructures and the cell surface in live Caulobacter crescentus,” Proc Natl Acad Sci U S A 108, E1102–1110 (2011).

4. F. Balzarotti, Y. Eilers, K. C. Gwosch, A. H. Gynna, V. Westphal, F. D. Stefani, J. Elf, and S. W. Hell, “Nanometer resolution imaging and tracking of fluorescent molecules with minimal photon fluxes,” Science 355, 606–612 (2017).

5. T. Deguchi, M. K. Iwanski, E.-M. Schentarra, C. Heidebrecht, L. Schmidt, J. Heck, T. Weihs, S. Schnorrenberg, P. Hoess, S. Liu, V. Chevyreva, K.-M. Noh, L. C. Kapitein, and J. Ries, “Direct observation of motor protein stepping in living cells using MINFLUX,” Science 379, 1010–1015 (2023).

6. K. H. Song, Y. Zhang, G. Wang, C. Sun, and H. F. Zhang, “Three-dimensional biplane spectroscopic single-molecule localization microscopy,” Optica 6, 709–715 (2019).

7. Y. I. Chen, Y. J. Chang, S. C. Liao, T. D. Nguyen, J. Yang, Y. A. Kuo, S. Hong, Y. L. Liu, H. G. Rylander, 3rd, S. R. Santacruz, T. E. Yankeelov, and H. C. Yeh, “Generative adversarial network enables rapid and robust fluorescence lifetime image analysis in live cells,” Commun Biol 5, 18 (2022).

8. M. Mazaheri, J. Ehrig, A. Shkarin, V. Zaburdaev, and V. Sandoghdar, “Ultrahigh-Speed Imaging of Rotational Diffusion on a Lipid Bilayer,” Nano Lett 20, 7213–7219 (2020).

9. D. Kim, Z. Y. Zhang, and K. Xu, “Spectrally Resolved Super-Resolution Microscopy Unveils Multipath Reaction Pathways of Single Spiropyran Molecules,” Journal of the American Chemical Society 139, 9447–9450 (2017).

10. L. Sansalone, Y. Zhang, M. M. A. Mazza, J. L. Davis, K. H. Song, B. Captain, H. F. Zhang, and F. M. Raymo, “High-Throughput Single-Molecule Spectroscopy Resolves the Conformational Isomers of BODIPY Chromophores,” Journal of Physical Chemistry Letters 10, 6807–6812 (2019).

11. W. Y. Kao, C. E. Davis, Y. I. Kim, and J. M. Beach, “Fluorescence emission spectral shift measurements of membrane potential in single cells,” Biophys J 81, 1163–1170 (2001).

12. A. S. Klymchenko, “Solvatochromic and Fluorogenic Dyes as Environment-Sensitive Probes: Design and Biological Applications,” Acc Chem Res 50, 366–375 (2017).

13. C. Phelps, T. Huang, J. Wang, and X. L. Nan, “Multipair Fo?rster Resonance Energy Transfer via Spectrally Resolved Single-Molecule Detection,” J Phys Chem B 126, 5765–5771 (2022).

14. K. Zhanghao, W. H. Liu, M. Q. Li, Z. H. Wu, X. Wang, X. Y. Chen, C. Y. Shan, H. Q. Wang, X. W. Chen, Q. H. Dai, P. Xi, and D. Y. Jin, “High-dimensional super-resolution imaging reveals heterogeneity and dynamics of subcellular lipid membranes,” Nature Communications 11 (2020).

15. M. N. Bongiovanni, J. Godet, M. H. Horrocks, L. Tosatto, A. R. Carr, D. C. Wirthensohn, R. T. Ranasinghe, J. E. Lee, A. Ponjavic, J. V. Fritz, C. M. Dobson, D. Klenerman, and S. F. Lee, “Multi-dimensional super-resolution imaging enables surface hydrophobicity mapping,” Nat Commun 7, 13544 (2016).

16. R. L. Zhang, F. W. Pratiwi, B. C. Chen, P. L. Chen, S. H. Wu, and C. Y. Mou, “Simultaneous Single-Particle Tracking and Dynamic pH Sensing Reveal Lysosome-Targetable Mesoporous Silica Nanoparticle Pathways,” Acs Appl Mater Inter 12, 42472–42484 (2020).

17. L. Sansalone, Y. Zhang, M. M. A. Mazza, J. L. Davis, K. H. Song, B. Captain, H. F. Zhang, and F. M. Raymo, “High-Throughput Single-Molecule Spectroscopy Resolves the Conformational Isomers of BODIPY Chromophores,” J Phys Chem Lett 10, 6807–6812 (2019).

18. R. L. Zhang, F. W. Pratiwi, B. C. Chen, P. Chen, S. H. Wu, and C. Y. Mou, “Simultaneous Single-Particle Tracking and Dynamic pH Sensing Reveal Lysosome-Targetable Mesoporous Silica Nanoparticle Pathways,” ACS Appl Mater Interfaces 12, 42472–42484 (2020).

19. J. E. Lee, J. C. Sang, M. Rodrigues, A. R. Carr, M. H. Horrocks, S. De, M. N. Bongiovanni, P. Flagmeier, C. M. Dobson, D. J. Wales, S. F. Lee, and D. Klenerman, “Mapping Surface Hydrophobicity of alpha-Synuclein Oligomers at the Nanoscale,” Nano Lett 18, 7494–7501 (2018).

20. B. Dong, L. Almassalha, B. E. Urban, T. Q. Nguyen, S. Khuon, T. L. Chew, V. Backman, C. Sun, and H. F. Zhang, “Super-resolution spectroscopic microscopy via photon localization,” Nat Commun 7, 12290 (2016).

21. Z. Zhang, S. J. Kenny, M. Hauser, W. Li, and K. Xu, “Ultrahigh-throughput single-molecule spectroscopy and spectrally resolved super-resolution microscopy,” Nat Methods 12, 935–938 (2015).

22. S. Moon, R. Yan, S. J. Kenny, Y. Shyu, L. Xiang, W. Li, and K. Xu, “Spectrally Resolved, Functional Super-Resolution Microscopy Reveals Nanoscale Compositional Heterogeneity in Live-Cell Membranes,” J Am Chem Soc 139, 10944–10947 (2017).

23. D. I. Danylchuk, S. Moon, K. Xu, and A. S. Klymchenko, “Switchable Solvatochromic Probes for Live-Cell Super-resolution Imaging of Plasma Membrane Organization,” Angew Chem Int Ed Engl 58, 14920–14924 (2019).

24. H. Guan, D. Li, H. C. Park, A. Li, Y. Yue, Y. A. Gau, M. J. Li, D. E. Bergles, H. Lu, and X. Li, “Deep-learning two-photon fiberscopy for video-rate brain imaging in freely-behaving mice,” Nat Commun 13, 1534 (2022).

25. N. Wagner, F. Beuttenmueller, N. Norlin, J. Gierten, J. C. Boffi, J. Wittbrodt, M. Weigert, L. Hufnagel, R. Prevedel, and A. Kreshuk, “Deep learning-enhanced light-field imaging with continuous validation,” Nat Methods 18, 557–563 (2021).

26. Y. Rivenson, Y. Wu, and A. Ozcan, “Deep learning in holography and coherent imaging,” Light Sci Appl 8, 85 (2019).

27. X. Huang, J. Fan, L. Li, H. Liu, R. Wu, Y. Wu, L. Wei, H. Mao, A. Lal, P. Xi, L. Tang, Y. Zhang, Y. Liu, S. Tan, and L. Chen, “Fast, long-term, super-resolution imaging with Hessian structured illumination microscopy,” Nat Biotechnol 36, 451–459 (2018).

28. S. Puntener, and P. Rivera-Fuentes, “Single-Molecule Peptide Identification Using Fluorescence Blinking Fingerprints,” J Am Chem Soc 145, 1441–1447 (2023).

29. Q. Wang, H. He, Q. Zhang, Z. Feng, J. Li, X. Chen, L. Liu, X. Wang, B. Ge, D. Yu, H. Ren, and F. Huang, “Deep-Learning-Assisted Single-Molecule Tracking on a Live Cell Membrane,” Anal Chem (2021).

30. Z. Wang, L. Zhu, H. Zhang, G. Li, C. Yi, Y. Li, Y. Yang, Y. Ding, M. Zhen, S. Gao, T. K. Hsiai, and P. Fei, “Real-time volumetric reconstruction of biological dynamics with light-field microscopy and deep learning,” Nat Methods 18, 551–556 (2021).

31. X. Chen, B. W. Li, S. W. Jiang, T. Zhang, X. Zhang, P. W. Qin, X. Yuan, Y. B. Zhang, G. A. Zheng, and X. Y. Ji, “Accelerated Phase Shifting for Structured Illumination Microscopy Based on Deep Learning,” Ieee T Comput Imag 7, 700–712 (2021).

32. A. Speiser, L. R. Muller, P. Hoess, U. Matti, C. J. Obara, W. R. Legant, A. Kreshuk, J. H. Macke, J. Ries, and S. C. Turaga, “Deep learning enables fast and dense single-molecule localization with high accuracy,” Nat Methods 18, 1082–1090 (2021).

33. Y. Zhang, G. Wang, P. Huang, E. Sun, J. Kweon, Q. Li, J. Zhe, L. L. Ying, and H. F. Zhang, “Minimizing Molecular Misidentification in Imaging Low-Abundance Protein Interactions Using Spectroscopic Single-Molecule Localization Microscopy,” Anal Chem 94, 13834–13841 (2022).

34. I. J. Goodfellow, J. Pouget-Abadie, M. Mirza, B. Xu, D. Warde-Farley, S. Ozair, A. Courville, and Y. Bengio, “Generative Adversarial Nets,” Adv Neur In 27, 2672–2680 (2014).

35. C. Ledig, L. Theis, F. Huszar, J. Caballero, A. Cunningham, A. Acosta, A. Aitken, A. Tejani, J. Totz, Z. H. Wang, and W. Z. Shi, “Photo-Realistic Single Image Super-Resolution Using a Generative Adversarial Network,” Proc Cvpr Ieee, 105–114 (2017).

36. T. C. Wang, M. Y. Liu, J. Y. Zhu, A. Tao, J. Kautz, and B. Catanzaro, “High-Resolution Image Synthesis and Semantic Manipulation with Conditional GANs,” 2018 Ieee/Cvf Conference on Computer Vision and Pattern Recognition (Cvpr), 8798–8807 (2018).

37. O. Ronneberger, P. Fischer, and T. Brox, “U-Net: Convolutional Networks for Biomedical Image Segmentation,” Lect Notes Comput Sc 9351, 234–241 (2015).

38. Y. Y. Lin, B. W. Zeng, Y. F. Wang, Y. Chen, Z. J. Fang, J. Zhang, X. Y. Ji, H. Q. Wang, and Y. B. Zhang, “Unpaired Multi-Domain Stain Transfer for Kidney Histopathological Images,” Aaai Conf Artif Inte, 1630–1637 (2022).

39. R. Kreder, K. A. Pyrshev, Z. Darwich, O. A. Kucherak, Y. Mely, and A. S. Klymchenko, “Solvatochromic Nile Red probes with FRET quencher reveal lipid order heterogeneity in living and apoptotic cells,” ACS Chem Biol 10, 1435–1442 (2015).

40. K. Dragomiretskiy, and D. Zosso, “Variational Mode Decomposition,” Ieee T Signal Proces 62, 531–544 (2014).

41. W. Teo, A. V. Caprariello, M. L. Morgan, A. Luchicchi, G. J. Schenk, J. T. Joseph, J. J. G. Geurts, and P. K. Stys, “Nile Red fluorescence spectroscopy reports early physicochemical changes in myelin with high sensitivity,” Proc Natl Acad Sci U S A 118 (2021).

42. J. A. Jackman, and N. J. Cho, “Supported Lipid Bilayer Formation: Beyond Vesicle Fusion,” Langmuir 36, 1387–1400 (2020).

43. S. Hou, J. Exell, and K. Welsher, “Real-time 3D single molecule tracking,” Nat Commun 11, 3607 (2020).

44. S. Hou, X. Lang, and K. Welsher, “Robust real-time 3D single-particle tracking using a dynamically moving laser spot,” Opt Lett 42, 2390–2393 (2017).

45. K. Xu, H. P. Babcock, and X. Zhuang, “Dual-objective STORM reveals three-dimensional filament organization in the actin cytoskeleton,” Nat Methods 9, 185–188 (2012).

46. M. Ovesny, P. Krizek, J. Borkovec, Z. Svindrych, and G. M. Hagen, “ThunderSTORM: a comprehensive ImageJ plug-in for PALM and STORM data analysis and super-resolution imaging,” Bioinformatics 30, 2389–2390 (2014).

47. J. Cheng, X. Chen, L. X. Xu, and Z. J. Wang, “Illumination Variation-Resistant Video-Based Heart Rate Measurement Using Joint Blind Source Separation and Ensemble Empirical Mode Decomposition,” Ieee J Biomed Health 21, 1422–1433 (2017).

48. K. M. He, X. Y. Zhang, S. Q. Ren, and J. Sun, “Deep Residual Learning for Image Recognition,” 2016 Ieee Conference on Computer Vision and Pattern Recognition (Cvpr), 770–778 (2016).

49. T. Miyato, T. Kataoka, M. Koyama, and Y. Yoshida, “Spectral Normalization for Generative Adversarial Networks,” presented at the International Conference on Learning Representations (ICLR) 2018.

50. I. Gulrajani, F. Ahmed, M. Arjovsky, V. Dumoulin, and A. C. Courville, “Improved Training of Wasserstein GANs,” presented at the Advances in Neural Information Processing Systems (NIPS) 2017.

