## Supplementary Information for "Deep learning-enhanced single-molecule spectrum imaging"

| Section | Page |
| --- | --- |
| **Figures** | |
| Figure S1: Optical setup of SR-SRM system | 1 |
| Figure S2: Overview of bicelle sample preparation | 2 |
| Figure S3: Calibration of pixel-wavelength curve | 3 |
| Figure S4: Workflow of data preprocess | 4 |
| Figure S5: Principle of VMD | 5 |
| Figure S6: Averaged spectra of 10 molecules in different solvents | 6 |
| Figure S7: Results comparison in SLBs with data screening | 7 |
| Figure S8: Averaged spectra of 10 molecules in SLBs with data screening | 8 |
| Figure S9: Centroid identification accuracy in SLBs with data screening | 9 |
| Figure S10: Averaged spectra of 10 molecules in SLBs with data with ResNet output | 10 |
| Figure S11: Centroid identification accuracy in SLBs with ResNet output | 11 |
| **Tables** | |
| Table S1: Pixel shifts measured under the corresponding bandpass filter | 12 |
| **Notes** | |
| Supplementary Note 1: Optical setup | 13 |
| Supplementary Note 2: Preparation for sample | 14 |
| Supplementary Note 3: Calibration of SR-SRM system | 16 |
| Supplementary Note 4: VMD algorithm | 17 |
| Supplementary Note 5: Network training schedule | 19 |

### Supporting Information

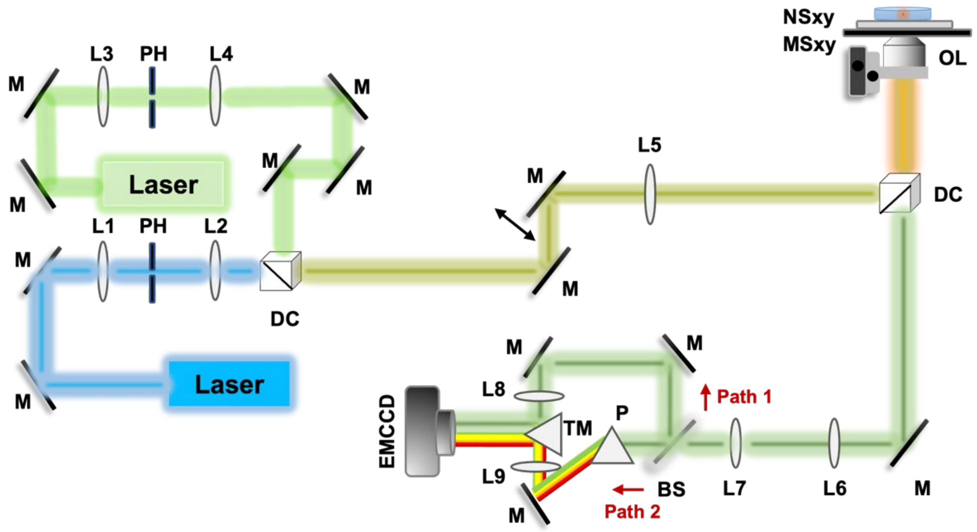


**Figure S1.** Optical setup. For the Nile Red-based spectrally resolved super resolution imaging, the sample was illuminated by a laser, and position and spectrum information of molecules were recorded simultaneously by the EMCCD.

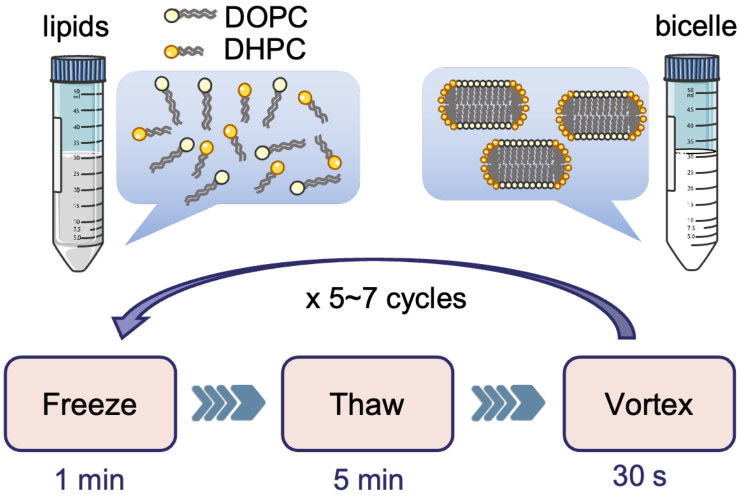


**Figure S2.** Overview of bicelle sample preparation.

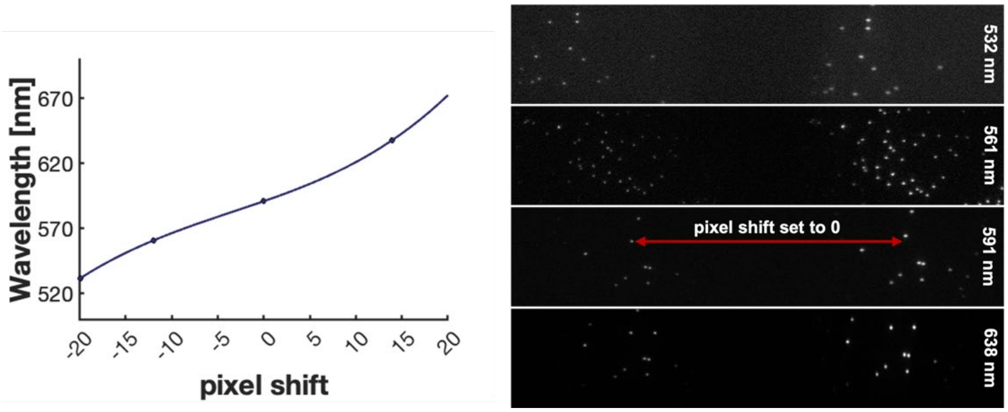


**Figure S3.** Calibration of pixel-wavelength curve. The pixel shift was mapping to the real wavelength value according to the pixel-wavelength curve (left). The paired data was collected using different bandpass filters (right).

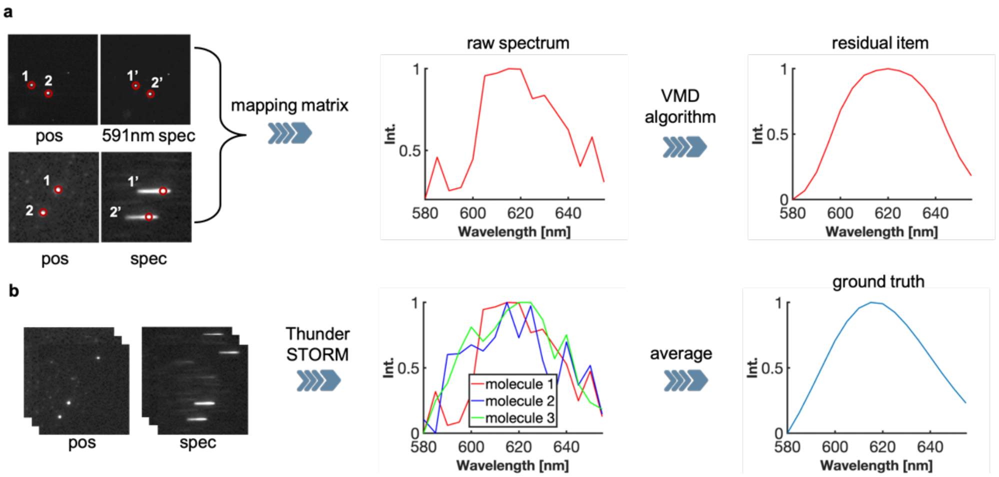


**Figure S4**. Workflow of data preprocess. (a) Data pre-processing process for a single frame. The residual terms of single-molecule spectra are extracted using the variational modal decomposition (VMD) algorithm. (b) Preparation of ground truth used for training. Each point coordinates of the molecules are determined by the ThunderSTORM ImageJ plugin.

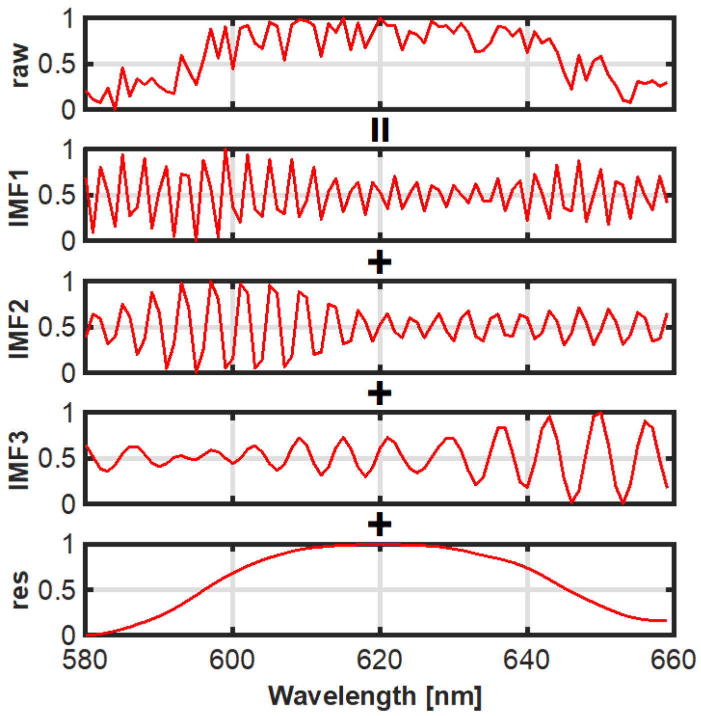


**Figure S5.** Principle of VMD. Different rows represent the corresponding components of the raw signal.

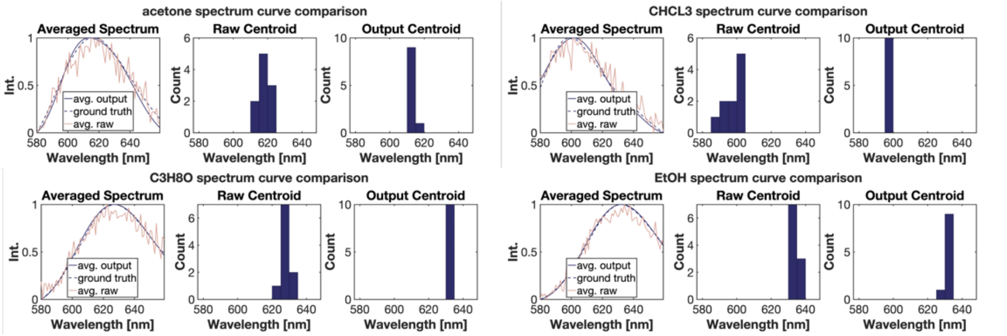


**Figure S6**. Averaged spectra of 10 molecules in different solvents. The real centroids of CHCl3, acetone, C3H8O, and EtOH are 600 nm, 617 nm 628 nm, and 632 nm, respectively.

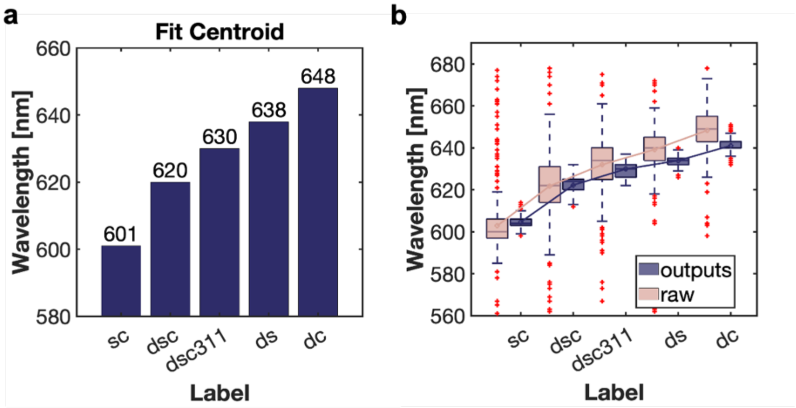


**Figure S7.** Results comparison in SLBs with data screening. (a) The fluorescence emission peak of Nile Red spectra in different components of SLBs. The real centroids of DC (DOPC:Chol=1:1), DS (DOPC:SM=1:1), DSC311 (DOPC:SM:Chol=3:1:1), DSC(DOPC:SM:Chol=1:1:1) and SC (SM:Chol=1:1) are 601 nm, 620 nm 630 nm, 638 nm, and 648 nm respectively. (b) Boxplot of centroid calculated by SpecGAN outputs after manually data screening.

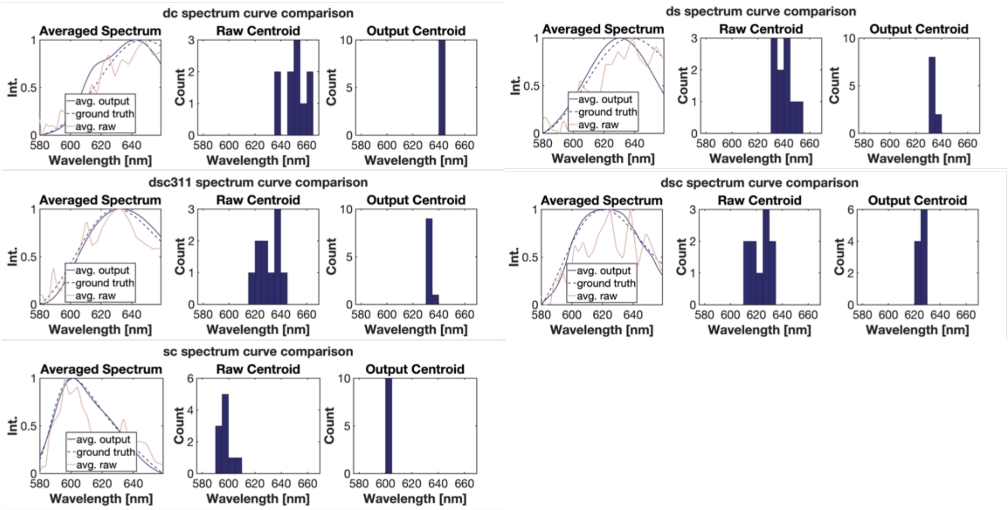


**Figure S8.** Averaged spectra of 10 molecules in SLBs with data screening. The cleaning method is to remove those data where the difference between the fitting centroid and the ground truth is greater than 15nm or the RMSE is greater than 0.3.

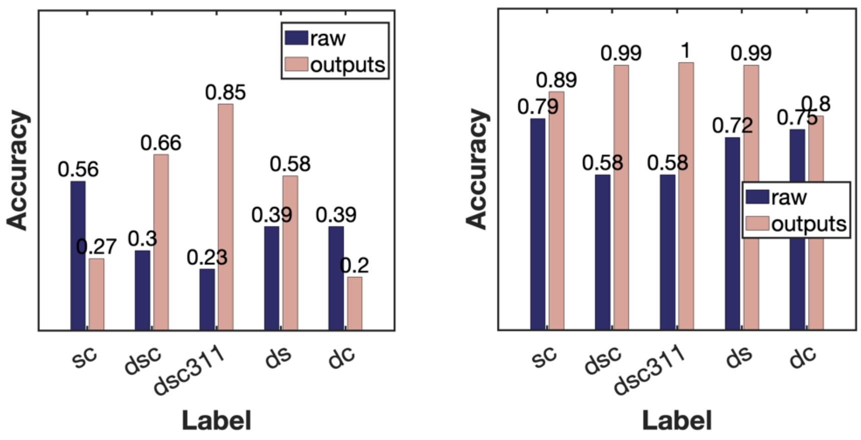


**Figure S9**. Centroid identification accuracy in SLBs with data screening. In the range of the average spectral centroid ±5 nm (left) and ±10 nm (right), the accuracy comparison of identifying the maximum emission peak position in the outputs of SpecGAN and raw data.

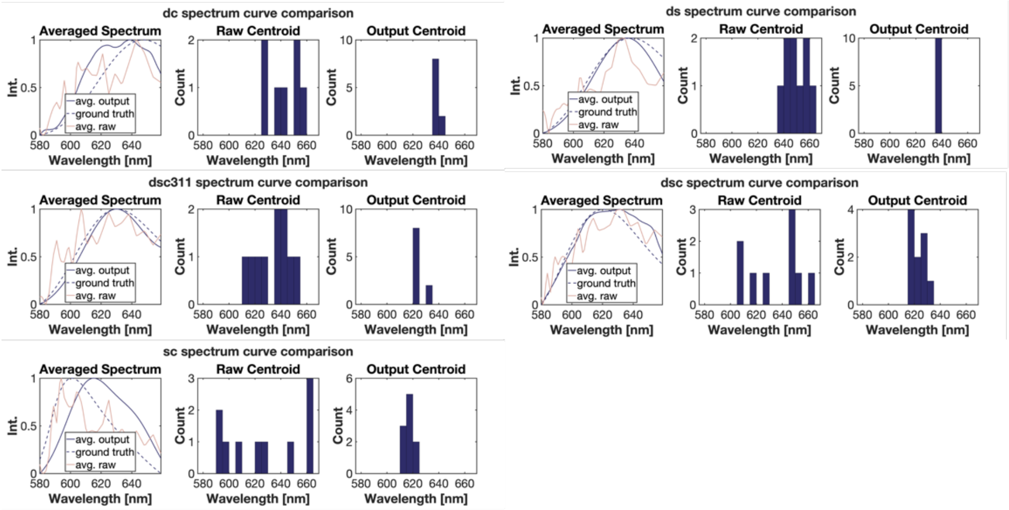


**Figure S10.** Averaged spectra of 10 molecules in SLBs with data with ResNet output. The ResNet-based classifier was used to determine the quality of signal, and SpecGAN only deal with those spectra being labeled as signal by classfier.

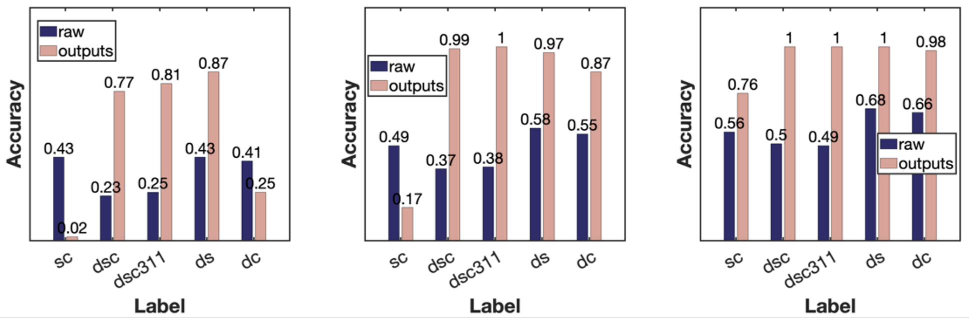


**Figure S11.** Centroid identification accuracy in SLBs with ResNet output. In the range of the average spectral centroid ±10 nm (left), ±15 nm (middle) and ±20 nm (right), the accuracy comparison of identifying the maximum emission peak position in the outputs of SpecGAN and raw data.

**Table S1.** Pixel shifts measured under the corresponding bandpass filter.

| wavelength | absolute pixel shifts | relative pixel shifts |
|---|---|---|
| 532 nm | 301.5 | -19.8 |
| 561 nm | 293.6 | -11.9 |
| 591 nm | 281.6 | 0 |
| 638 nm | 267.7 | 13.9 |

#### Supplementary Note 1: Optical setup

We manually built a SR-SRM system based on total internal reflection inverted fluorescence (TIRF) microscope, as shown in Fig S1. The 488-nm (LBX-488-60-CSB-PPA, OXXIUS) and 561-nm (LCX-561L-50-CSB-PPA, OXXIUS) lasers were manipulated to obtain desired sizes and shapes of spots by using relay lenses (L1/L3, $\boldsymbol{f}$=100mm; L2/L4, $\boldsymbol{f}$=200mm) and pinholes positioned in the focal plane of L1/L3 with a size of 100 μm. By changing the position of TIRF mirror via a one-dimensional translation stage, the angle of incident beams impinges on the sample was able to be modified. If the incident angle was equal to the critical angle, the total internal reflection illumination was achieved [1]. A TIRF lens (L5, $\boldsymbol{f}$=250 mm) was placed at the front focal plane of the 100× oil-immersion objective lens (Olympus, UPLAPO100XOHR, NA 1.5) to achieve wide-field illumination. When the fluorescent molecule was excited by laser, the emission signal reaches the imaging path through the dichroic mirror (ZT488/561rpc-UF1, Chroma). In the imaging path, the emission fluorescence was split into two paths by a 30:70 beam splitter (BSS10R, Thorlabs). In both path1 and path2, the information of molecules was focused by achromatic lens (L8 and L9, $\boldsymbol{f}$=150 mm), thus the effective magnification was ~83×. In path2, the fluorescent signal from the molecule was dispersed by an equilateral calcium fluoride (CaF2) prism (PS863, Thorlabs), with the spectral resolution being dependent on the distance between prism and the EMCCD camera (iXon 897, Andor). By combing the position and spectrum information using a right-angle prism mirror (MRA25L-E02, Thorlabs), the EMCCD (iXon 897, Andor) can concurrently record the positions and spectrum of molecules with a frame size of 512×256 pixels. For the SR-PAINT or SR-STORM imaging, the 561-nm laser power was set to 20 mW, and a sequence of frames was recorded with 20 ms exposure time and 200 gains until the fluorescent molecules were bleached. To exclude the interference of laser reflection, three different filters (ET570lp, Chroma, ET575lp, Chroma, and FF01-630/92, Semrock) were employed in the SR-STORM or SR-PAINT system.

#### Supplementary Note 2: Preparation for sample

##### Supported lipid bilayers (SLBs)

Supported lipid bilayers (SLBs) are model membranes that are widely used in the study of biological membranes. The lipid bilayer consists of two leaflets of phospholipid molecules with hydrophilic head groups facing the aqueous solution and hydrophobic tails facing each other. The hydrophobic tails form a fluid and dynamic interior region, which acts as a barrier to hydrophilic molecules, while the hydrophilic head groups interact with water and other polar molecules. SLBs are often used as model systems to study the properties and behaviors of cell membranes, such as membrane dynamics, protein-lipid interactions, and signaling pathways. For the preparation of SLBs, there are several methods to form SLBs including extrusion method, vesicle fusion method and bicelle method. Here we adopted the bicelle method to form stable SLBs on the surface of coverslip.

The details of bicelle sample preparation is reported before [2], and the overview is depicted in **Fig.S2**. Briefly, Sphingomyelin, cholesterol, and 1,2-dihexanoyl-sn-glycero-3-phosphocholine (DHPC) were purchased from Aladdin (S130559, C104028, D130408). 1,2-dioleoyl-sn-glycero-3-phosphocholine (DOPC) was purchased from Macklin (D838350). Each lipid component was dissolved in chloroform to 10mg/mL separately and then stored in -20℃ protecting from light. To prepare SLBs, the chloroform solutions of DOPC, sphingomyelin (SM), and cholesterol (Chol) were mixed in a 50 mL EP centrifuge tube with a certain ratio. After evaporation of chloroform from the long-chain lipid (DOPC mixture) and the short-chain lipid (DHPC) solution with nitrogen flow, the centrifuge tube was placed in a vacuum desiccator overnight to remove the residual chloroform. The long-chain and short-chain lipid was next mixtured in 10 mL buffer (10 mM Tris-HCl, pH=7.4 and 150 mM NaCl), and the ratio of long-chain phospholipids to short-chain phospholipids was 0.25. The obtained lipid suspension had a DOPC mixture concentration of 1 mM and a DHPC concentration of 0.25 mM. The lipid suspension was frozen in liquid nitrogen for ~1 min, then heated in 60 °C water bath for ~5 min, and vortexed for ~30 s. After around 5−7 cycles per sample, the resulting suspension is visually clear at room temperature, and the bicelle suspend formed.

Before incubation, it is necessary to dilute the bicelle suspension with buffer to a concentration of 0.031 mM, which is approximately 30 times. Subsequently, the coverslip is incubated in the diluted bicelle suspension at 60 °C for 15 minutes. Finally, the buffer was replaced by Dulbecco's phosphate-buffered saline (DPBS) for imaging.

##### Nile Red staining

Nile Red is an uncharged hydrophobic molecule and is a lipid fluorescent dye. The 10mg Nile Red were dissolved in 10.47 mL DMSO solution to form a 3 mM stock solution. The stock solution was then diluted with DPBS to 100 nM and dispensed into 1.5 mL centrifuge tubes, stored in -80℃ protecting from light. For the SR-STORM imaging of fixed cells, cells were incubated with 100 nM Nile Red for 30 min, and then rinsed two or three times by DPBS. For the SR-PAINT imaging of fixed cells or SLBs, the Nile Red is further dissolved to 3nM with the DPBS containing 100-200 uM ascorbic acid (Sigma-Aldrich, PHR1008) to reduce the effect of fluorescent bleaching.

**Cell culture**

In cellular data, the COS-7 cells were cultured in Dulbecco's Modified Eagle's Medium (DMEM) with 9% fetal bovine serum (FBS) and 1% penicillin/streptomycin for 2 days in 5% $CO_2$ at 37°C. Before imaging, the cells were treated with 3% paraformaldehyde and 0.1% glutaraldehyde in DPBS for 20 min, followed by a 5 min rinse with DPBS containing 0.1% NaBH4 solution, and three washes with DPBS. In order to reduce the effect of fluorescent bleaching, the Nile Red is dissolved in the DPBS containing 100-200 uM ascorbic acid to 3 nM. The parameters of EM-CCD were the same as those in SLBs imaging but record 12,000 frames to achieve the spectrally super-resolution imaging of the fixed cells.

**Supplementary Note 3: Calibration of SR-SRM system**

We used the SR-SRM shown in **Fig.S1** to perform multispectral imaging of two types of fluorescent spheres (FSDG002 480/520, FSSY002 540/640, Bangs Lab). The pixel shifts corresponding to special wavelength were determined via different narrow bandpass filters The pixel shift corresponding to 591-nm was set to 0, and the mapping relationship between pixel shifts and wavelength was fitted with a third-order polynomial curve, as depicted in **Fig.S3**.

In 591-nm channel, at least 6 position-spectrum paired points were select to calculate to position-spectrum coordinate transformation matrix. Therefore, the construction of the raw spectrum of single molecule requires the position-spectrum coordinate transformation matrix and pixel shift-wavelength curve. The preparation of dataset is shown in **Fig.S4**, the dye molecule coordinate position detection was performed by ThunderSTORM, a modular plug-in for ImageJ designed for data processing of single-molecule localization microscopes. To improve the quality of raw signal, these spectra were further cleaning according to some statistical characteristics, such as coefficient of variation, average spectral photon number, kurtosis, etc. The input of SpecGAN is the residual item of raw spectrum, more details are depicted in the introduction of VMD algorithm. Averaging all spectra in the same environment is able to obtain the ground truth of spectrum of Nile Red.

**Supplementary Note 4: VMD algorithm**

In 2014, Dragomiretskiy and Zosso developed a variational mode decomposition (VMD) method, which estimates each signal component by solving the frequency domain variational optimization problem [3]. The raw signal can be decomposed to a series of intrinsic mode functions (IMFs), as shown in **Fig.S5**, and denoted as:

$$u_k(t) = a_k(t)\cos\big(\phi_k(t)\big), \tag{1}$$

where $\phi_k(t)$ is a non-decreasing function, which is $\phi'_k(t) \geq 0$, and the signal envelop $a_k(t)$ is non-negative. For a real signal $s_k(t)$, its analytic signal $s(t)$ can be written as:

$$s(t) = u_k(t) + jH[u_k(t)] = a_k(t)e^{j\phi(t)}, \tag{2}$$

where $H$ represents the Hilbert transform. The introduction of the analytic signal allows the lossless preservation of the positive frequency components and also increases the representability of the signal. The VMD algorithm assumes that all components are narrow-band signals concentrated around their respective center frequencies, thus the optimization problem is constructed as:

$$\min_{\{\mu_k,\omega_k\}}\left\{\sum_k \left\|\partial_t\left[\left(\delta(t)+\frac{j}{\pi t}\right)*u_k(t)\right]e^{-j\omega_k t}\right\|_2^2\right\} \quad s.t. \sum_k u_k(t) = f\ , \tag{3}$$

where $\omega_k$ corresponds to the center frequency of the k-th component. $\delta(t)$ is the Dirac function, and * denotes the operation of convolution.

To reconstruct the constraint problems, VMD algorithm introduce the augmented Lagrangian as follows:

$$\begin{aligned}\mathcal{L}(\{u_k\},\{\omega_k\},\lambda) := \alpha\sum_k \left\|\partial_t\left[\left(\delta(t)+\frac{j}{\pi t}\right)*u_k(t)\right]e^{-j\omega_k t}\right\|_2^2 \\ +\left\|f(t)-\sum_k u_k(t)\right\|_2^2 + < \lambda(t), f(t)-\sum_k u_k(t) > \end{aligned}, \tag{4}$$

where $f(t)$ is the raw signal and $\lambda$ represents the Lagrangian multipliers. The solutions to the constraint problem of **eq.(4)** can be achieved via the alternating direction method of multipliers (ADMM) method. Briefly, the idea is to fix two other variables and updating one of them, as demonstrated bellow:

$$\begin{aligned}u_k^{n+1} = argmin_{u_k \in X}\{\alpha\sum_k \left\|\partial_t\left[\left(\delta(t)+\frac{j}{\pi t}\right)*u_k(t)\right]e^{-j\omega_k t}\right\|_2^2 \\ +\left\|f(t)-\sum_k u_k(t)+\frac{\lambda(t)}{2}\right\|_2^2\}\end{aligned}. \tag{5}$$

Because of the property of the Parseval Fourier isometry under the L2 norm, this problem can be solved in spectral domain:

$$u_k^{n+1} = argmin_{u_k \in X}\{\int_0^{\infty} 4\alpha(\omega-\omega_k)^2 |\mu_k(\omega)|^2 + 2\left|f(\omega) - \sum_i u_i(\omega) + \frac{\lambda(\omega)}{2}\right|^2 d\omega\} . \quad (6)$$

Derivation of the above equation shows the solution of this quadratic optimization problem:

$$u_k^{n+1} = \frac{f(\omega) - \sum_{i \neq k} u_i(\omega) + \frac{\lambda(\omega)}{2}}{1 + 2\alpha(\omega-\omega_k)^2}. \quad (7)$$

For the update of $\omega_k^{n+1}$, the relevant problem has the format as shown:

$$\omega_k^{n+1} = argmin_{\omega_k}\left\{\left\|\partial_t\left[\left(\delta(t) + \frac{j}{\pi t}\right) * u_k(t)\right] e^{-j\omega_k t}\right\|_2^2\right\}. \quad (8)$$

We also deal with this optimization problem in Fourier domain, thus the **eq.(8)** can be denoted as:

$$\omega_k^{n+1} = argmin_{\omega_k}\{\int_0^{\infty} (\omega-\omega_k)^2 |\mu_k(\omega)|^2 d\omega\}, \quad (9)$$

by the same token, the update of $\omega_k^{n+1}$ is:

$$\omega_k^{n+1}(\omega) = \frac{\int_0^{\infty} \omega |u_k(\omega)| dw}{\int_0^{\infty} |u_k(\omega)| dw}, \quad (10)$$

**Supplementary Note 5: Network training schedule**

We used a deep learning server with 4 NIDIA GeForce RTX 2080Ti GPUs for model training and testing, and the code framework was based on Pytorch. For the training stage, the parameters of generator and discriminator were updated alternatively via the adaptive moment estimation (Adam) optimizer. The initialization of weights was follows a kaming uniform distribution. During training, the initial learning was $1\times10^{-4}$ and decreased with a variable decay. For more details of our SpecGAN model, it can be found on our github homepage.

### References


[1] D. S. Johnson, J. K. Jaiswal, and S. Simon, "Total internal reflection fluorescence (TIRF) microscopy illuminator for improved imaging of cell surface events," *Curr Protoc Cytom,* vol. Chapter 12, p. Unit 12 29, Jul 2012, doi: 10.1002/0471142956.cy1229s61.

[2] J. A. Jackman and N. J. Cho, "Supported Lipid Bilayer Formation: Beyond Vesicle Fusion," *Langmuir,* vol. 36, no. 6, pp. 1387-1400, Feb 18 2020, doi: 10.1021/acs.langmuir.9b03706.

[3] K. Dragomiretskiy and D. Zosso, "Variational Mode Decomposition," *IEEE Transactions on Signal Processing,* vol. 62, no. 3, pp. 531-544, 2014-02-01 2014, doi: 10.1109/tsp.2013.2288675.
